# RAPD amplicon sequence based metagenomics analysis of microbial communities associated with clove (*Syzygium aromaticum*) from Kerala, India

**DOI:** 10.1101/2023.03.14.532569

**Authors:** Sreekala Gangappan Santhakumari, Ramachandran Sarojini Santhosh

## Abstract

Four clove clones (BRC1, BRC3 MRC5 and MRC7) were collected from different places of ponmudi hills, Kerala, India. DNA was extracted and RAPD analysis was designed to evaluate clove clonal variation. Upon analysis bacterial sequences were also identified among clove genome sequences. So the clove and microbial sequences were separated and performed an analysis to identify clove associated metagenome. The major microbes present were *Mycobacterium canettii Xanthomaonas euvesicateria, Klebsiella* and *Psuedomonas*. A total of 88 genus of microbes were present altogether in four clones of *Syzygium aromaticum*. There were few fungal species were also identified, *Fusarium pseudograminearum* was the major fungus associated with all four different clones of clove.

## Introduction

Biome associated with plant and animals have role in their growth and development and also have a role in protecting from other microbes which may cause disease to them thus it may be beneficial or pathogenic. In this study, we concentrated on identifying microbes associated with plants using RAPD as a case study. Detailed knowledge of the microbial diversity, abundance, composition, functional genes patterns, and metabolic pathways at genome level could assist in understanding the contributions of microbial community towards plant growth and health, of plant-associated microbiota and plant metagenome, which includes bacteria, archaea, fungi, viruses and oomycetes. The nonpathogenic association may be mutualistic, beneficial, commensal or neutral. Prominent among organisms involved in beneficial association with the plant is endophyte. A plant contains different nutrients which aid in attracting microorganisms to it. Microorganisms found on the surface of the plant are called epiphytes while those who inhabit plant tissues are called endophytes. Plants present three types of environments depending on the one found to be conducive for the microbiota. The first environment is called the rhizosphere, this is where microorganism interacts with the soil and roots, the environment also contains many exudates from the plant. Endosphere is the second environment and its means inside the tissues of the plant. The third environment is called the phyllosphere which comprises the surface of the leaves and stems. The phyllosphere is known not to be a conducive environment for microorganisms, this because, nutrients availability in this environment is limited, irradiation of the sun is strong and water availability is inconsistent. Plant-associated microorganisms are regarded as mutualist, commensals or pathogens, the pathogenic ones are of great concern to scientist because of their economic importance. This study also focused on the mutualist found in the endosphere e.g. endophytes, which by definition, cause no harm to the plant. It also important to state that under certain conditions, plants also benefit from the numerous genes and proteins present in these microbes. Plant associated metagenome can improve our understanding of the importance of microbes to plants and the associations that exist among them. Presently, many public genomic databases for plant-associated microbial communities and metagenome are developed each year. DNA extraction from the plant tissues is important in plant metagenome studies. Diversity of endophytes in plants were recently studied. Plants endophytes are the most popular among the beneficial ones. These microorganisms make up a complex community and establish an intimate relationship with the host which researchers are just getting to understand in recent years.

## Results

### 16S metagenome

Among the major phylum actinobacteria, proteobacteria, firmicutes, bacteriodata, cyanobacteria, aquificae, and deinino coccus. Deinociccus were absent in MRC5 Aquificae were present only in BRC1, spirochaetes were present only in BRC3. Maximum number of actinobacteria and less diversity were observed in MRC5. Higher diversity was observed in BRC3.

**Fig.1.**
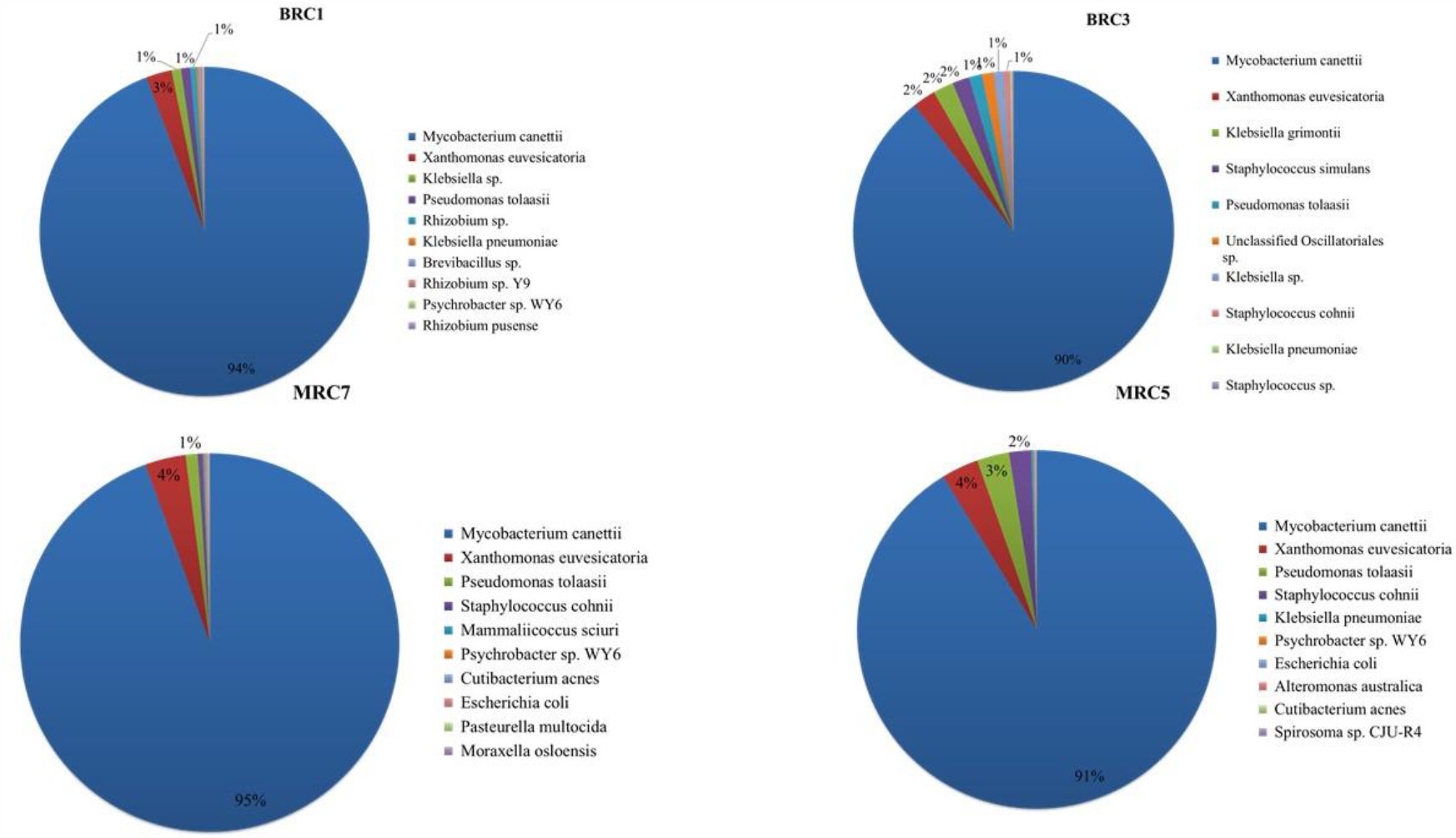
The istribution major microbial species present in four different clones of Syzygium aromaticum

Major number of actinomycetia were observed in MRC5 followed by BRC1, BRC3 and MRC7. Bacilli were also major in MRC5 followed by BRC3, MRC7 and BRC1, gammaproteobacteria were also major in MRC5 followed by BRC3, BRC1 and MRC7. Acidimicrobia, Aquificae bacteriodata, were present only in BRC1. Spirochaetia were present online BRC3.

Only present in MRC5 aeromonadales and streptomycetales, BRC1 acidimicrobiales, bacteroidales, eubacteriales MRC7-Neisseriales, rhodobacterales, Corynebacteriales were highest in MRC5 followed by BRC1, BRC3 and MRC7. Bacillales were higher in MRC5 followed by BRC3, MRC7 and BRC1. Eubacteriles were higher in BRC3, followed by BRC1, MRC5 and MRC7. Hyphomicrobiales were higher in BRC1. Oscillatoriles were higher in BRC3. Psuedomonadales and Xanthomonadales were higher in MRC5.

were major in all four clones of plant followed by firmicutes, proteobacteria and bacteroidota. Actinomycetia, gammaproteobacteria, deltaproteobacteria, cyanobacteria were present only in BRC3. Bacteria belonging to bacillales were major followed by acidomicrobials, xanthomonadales, spirochaetales and psuedomonadales Mycobacteriaceae family was major followed by Alteromonas australica was higher in MRC5. Brevibacillus sp. was higher in BRC1, Escherichia coli Was high in MRC5, Klebsiella grimontii was present only in BRC3, only Klebsiella pneumoniae were present in MRC5 and 7, Limnospira indica in BRC3 only, Mammaliicoccus sciuri was in MRC7 only, Mycobacterium canettii was present in all in higher amount MRC5, BRC1, BRC3 and MRC7.

Pseudomonas tolaasii Xanthomonas euvesicatoria and Staphylococcus cohnii were higher in MRC5, Staphylococcus simulans Were present in BRC3 only.

Species richness was high in BRc1 followed by MRC7, BRC3 and MRC5 The PCA plot shows the diversity between BRC1 and BRC3 are less but in between MRC and & there is a large diversity.

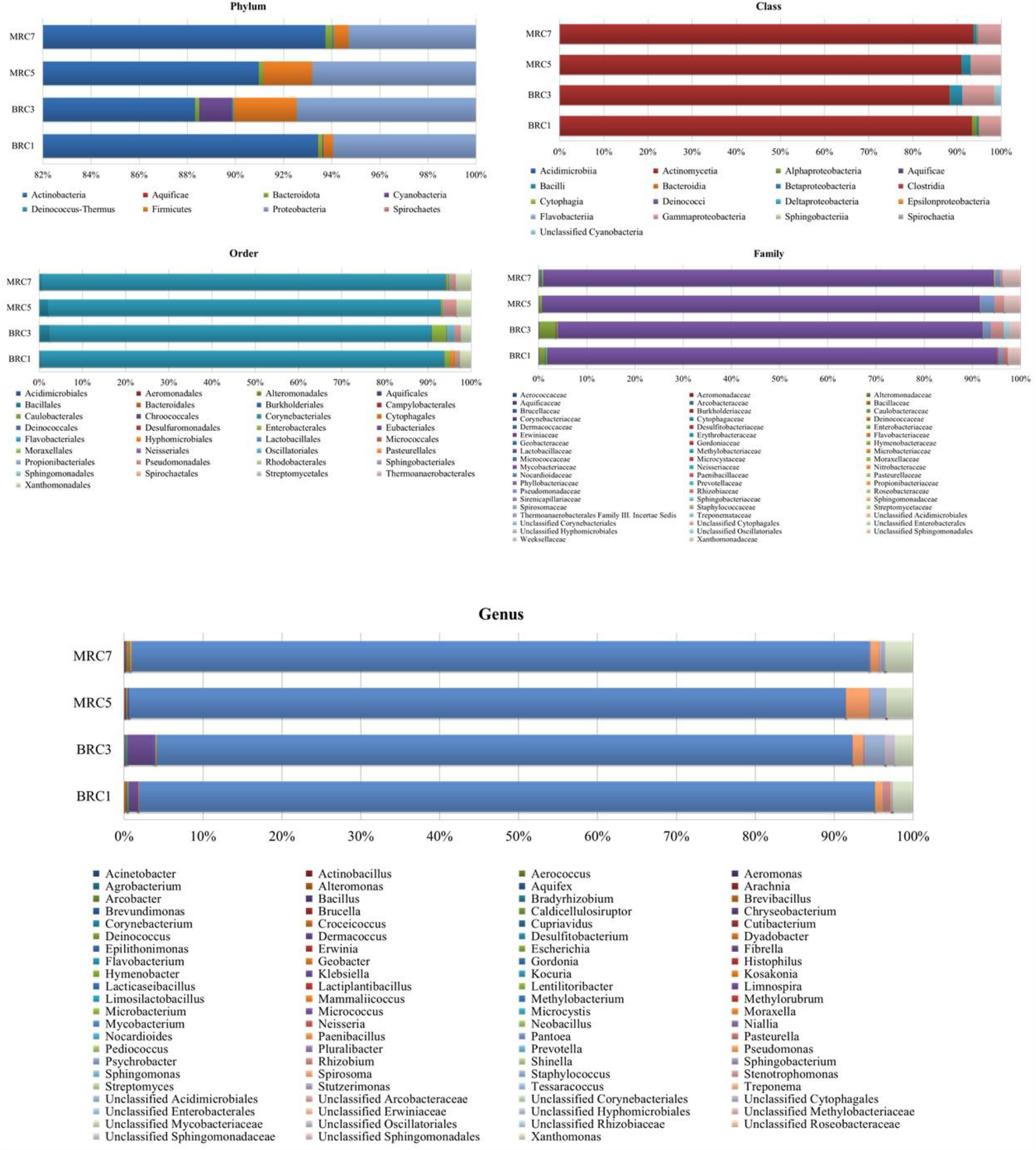

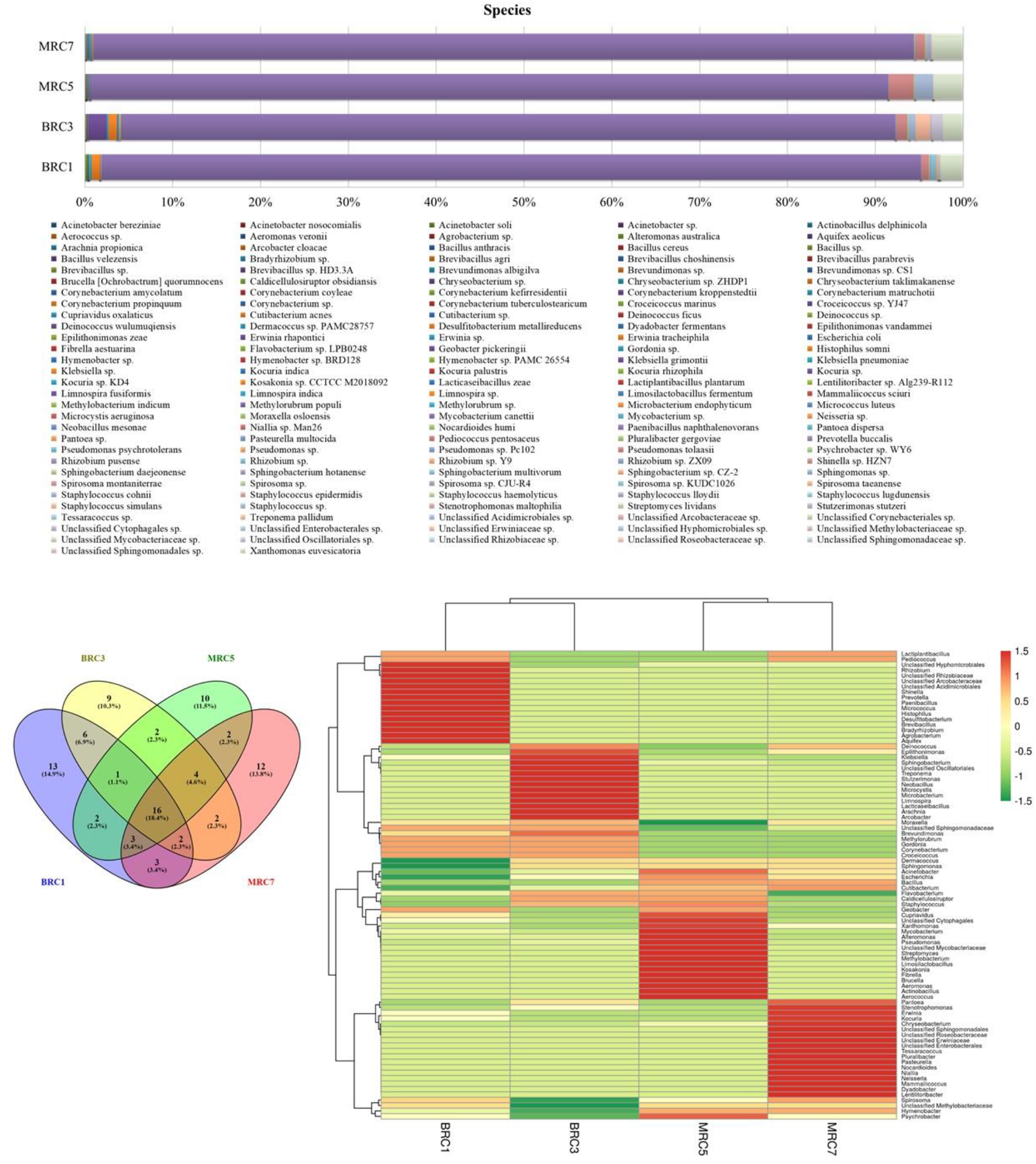

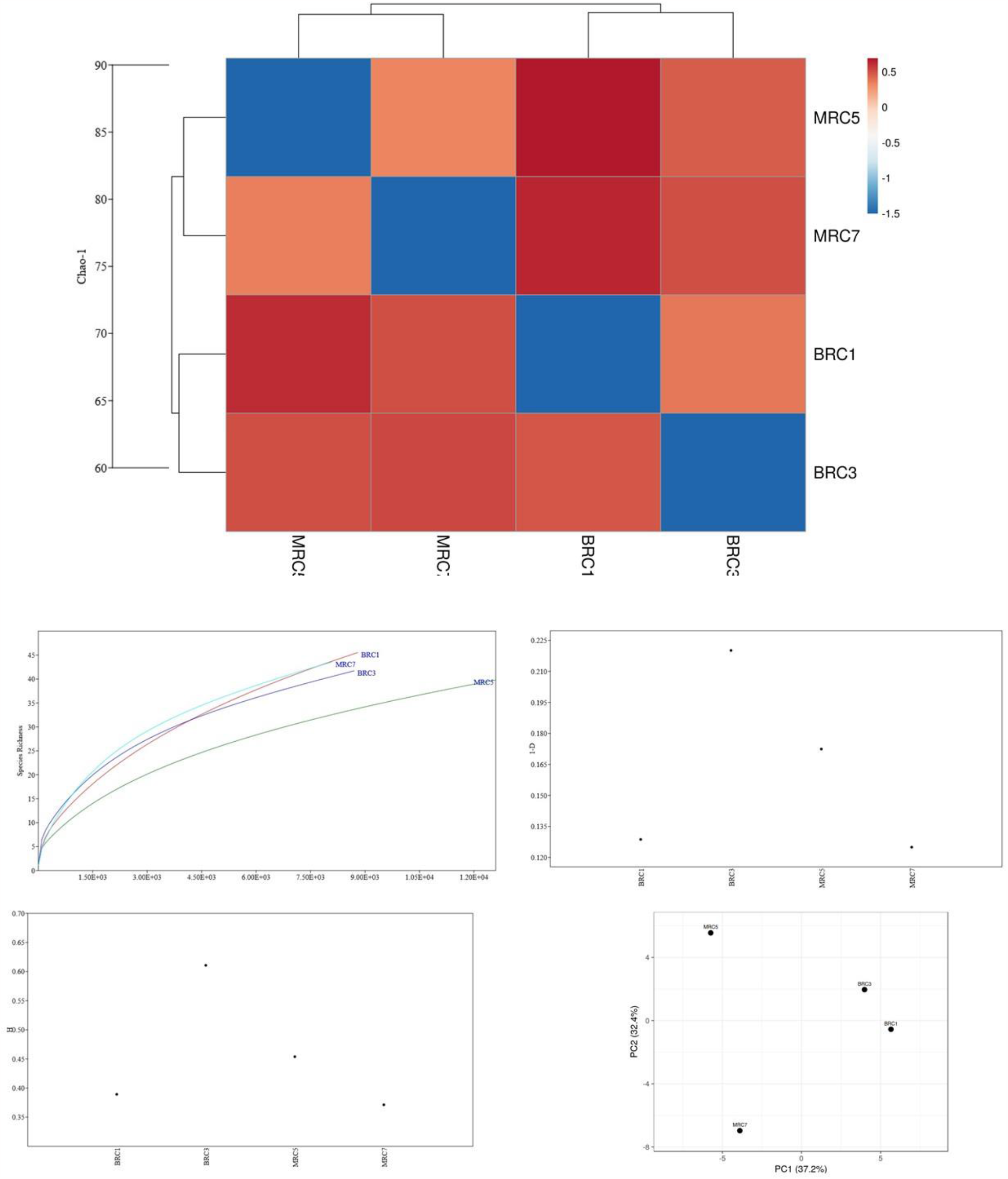

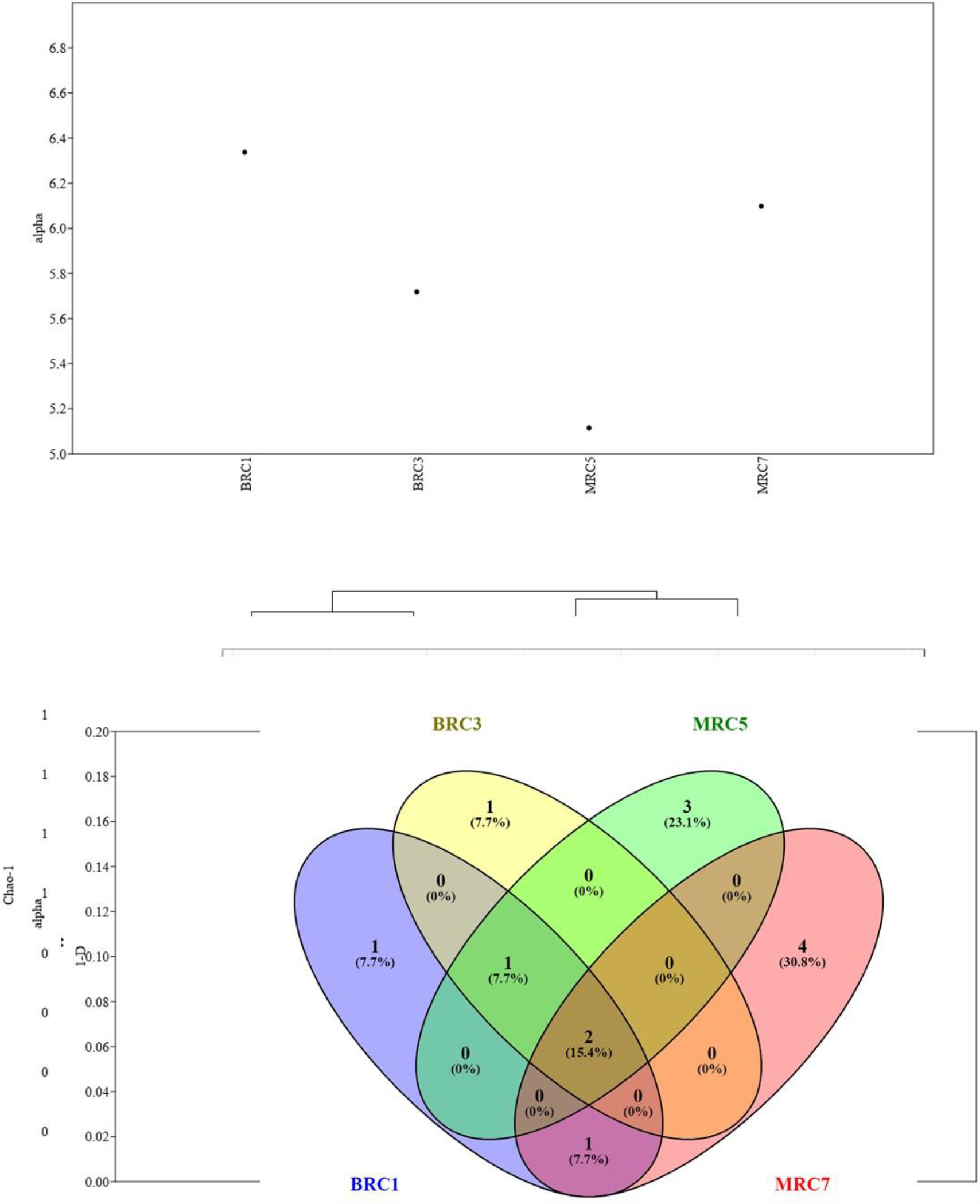

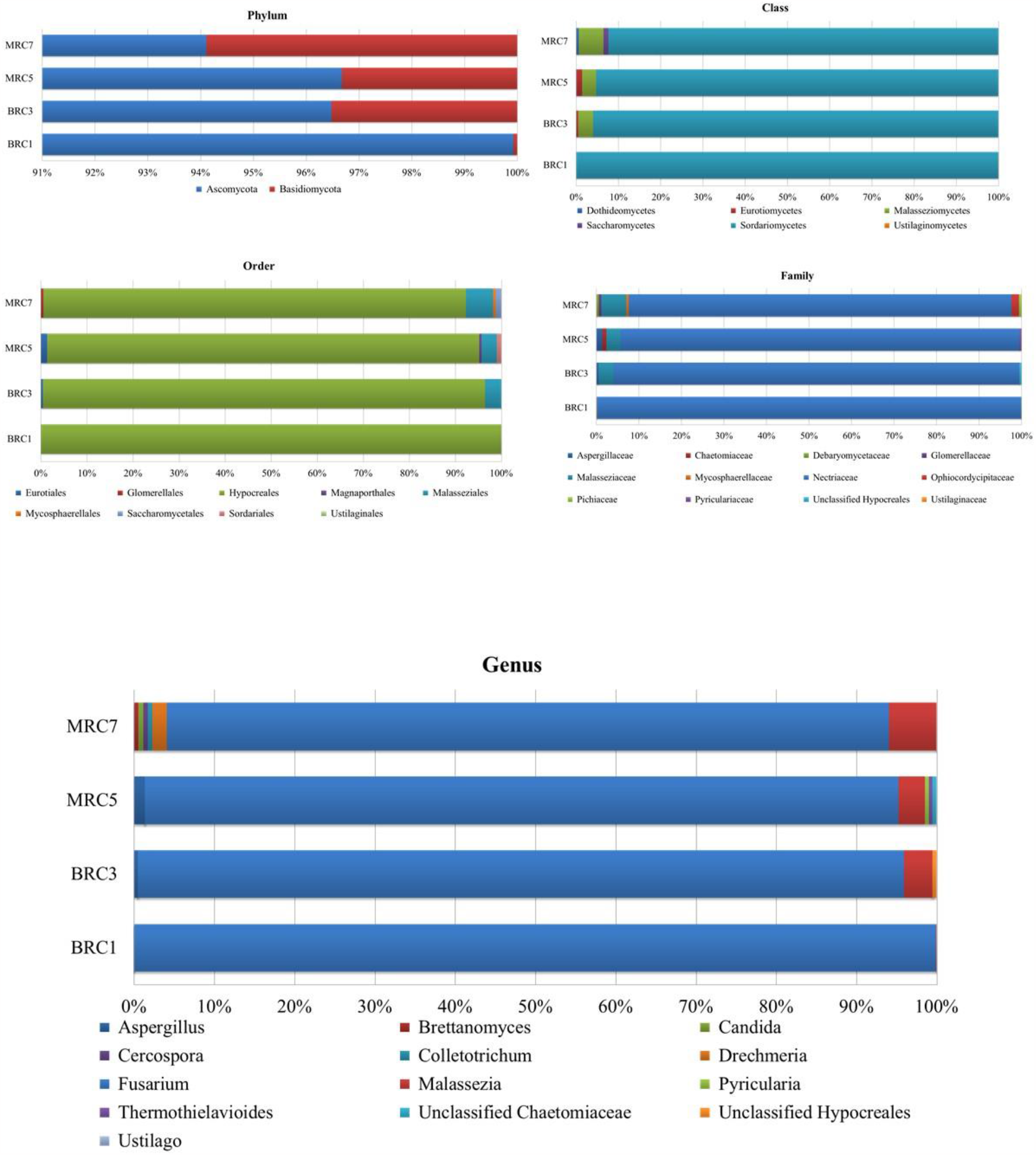

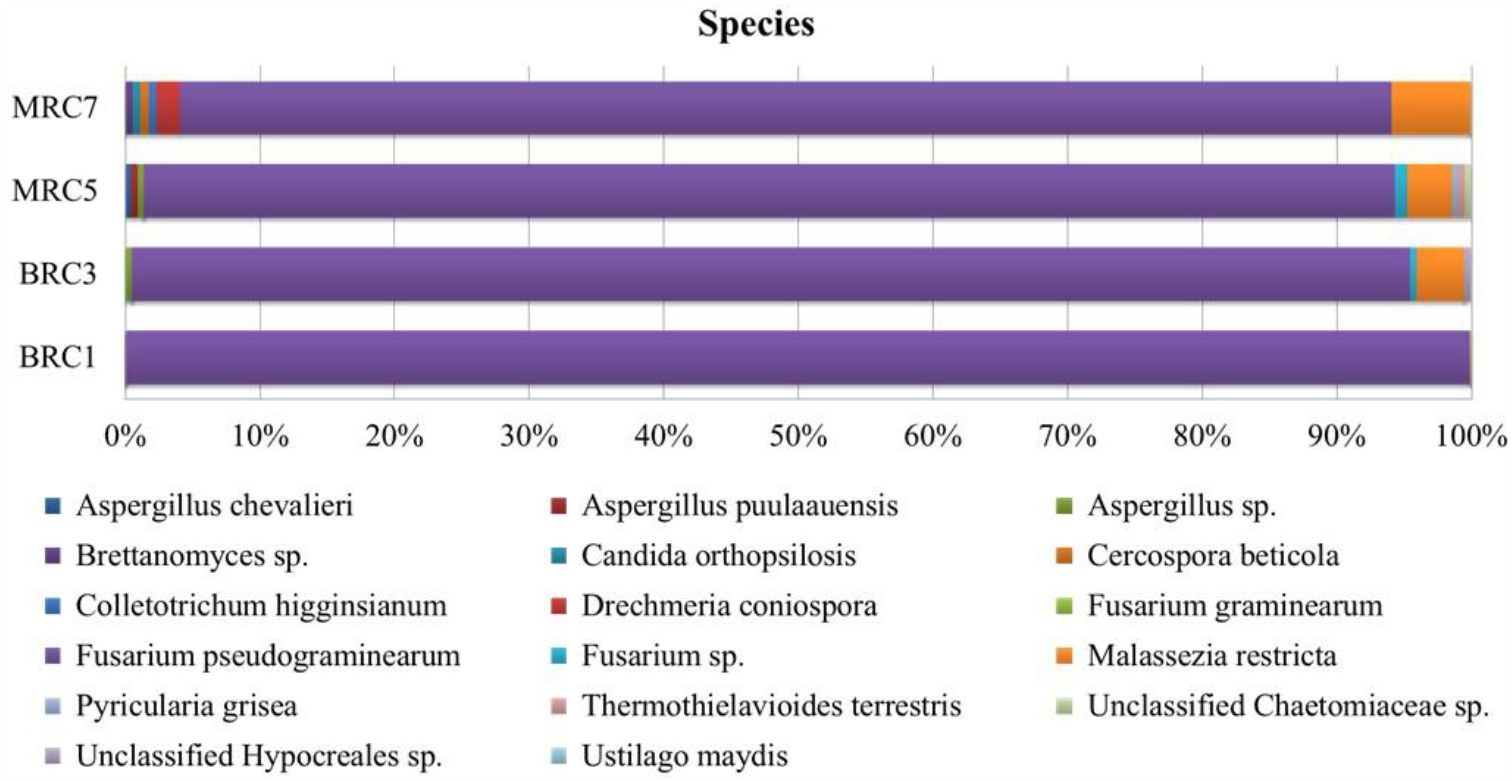

## Materials and Methods

### Sample Collection

#### Genomic DNA isolation

5g buds were kept at -70°C along with mortar and pestle for overnight. The buds were ground along with 500 mg Poly vinyl pyrollidone to powder form. The contents were transferred to a centrifuge tube and suspended in 8 ml suspension buffer (50mM EDTA, 120mMTris Hcl, 1M NaCl,0.5M sucrose, 2% Triton-X100, 0.2% beta-mercaptoethanol), incubated at 60°C for 30 min. After that 6 ml extraction buffer (20mM EDTA, 100mMTris Hcl, 1.5M NaCl, 2%CTAB, 1% beta-mercaptoethanol) was mixed and again incubated at 60°C for 30 min. The content was centrifuged at 12000 rpm for 15 min. at room temperature. The aqueous phase was collected to a new tube and added double volume of chloroform : isoamylalcohol mix. After incubation for 15 min. this was then centrifuged at 12000 rpm for 15 min. The aqueous phase was collected to a new tube and added 2v of chilled isopropanol and incubated at -20°C for 1h. The supernatant was discarded after centrifuged at 12000 rpm for 15 min. The pellet was re-suspended in chilled 70% ethanol, again centrifuged at 12000 rpm for 15 min. The supernatant was discarded and air dried. 500 μl sterile MQ water was then added for resuspension of DNA.

### 18SrRNA amplification

For PCR 1 ng of genomic DNA was used as template. Universal ITS4 and ITS5 sequence were used as forward and reverse primer sequences. Annealing temperature was 53°C for 30 sec. with 40 cycles. The PCR products were purified and sequenced by sanger’s method.

### Polymerase Chain Reaction

For obtaining RAPD fragments PCR was performed using 1 ng of genomic DNA as template. OPB1 to OPB10 primer were used. PCR was done as 94°C - 1 minute, 94°C - 30 seconds, 37°C – 1 minute, 68° C - 2 minutes, 68° C - 10 minutes, 4°C - ∞.. with 40 cycles. The PCR products were analysed in a 1% agarose gel.

### Gel analysis

The images were marked for polymorphic bands and table containing dominant marker of each plant were tabulated. Based on the table PIC value were calculated and based on that a phylogenetic tree was constructed.

### Sequencing of PCR fragments

From each plant all RAPD fragments were pooled from reactions OPB1 to OPB10. All four samples were subjected to NGS illumina sequencing using Analyser.

### Sequence analysis

#### Statistical Analysis

Statistical analyses were carried out in R (4.0.3). *vegan* (v 2.5-6) was used for all alpha-diversity calculations: Shannon diversity index (alpha diversity measurement of evenness and richness); evenness (homogeneous the distribution of taxa counts) and richness (number of taxa in a community). Pairwise Bray-Curtis dissimilarity was used to assess beta-diversity, or the overall variation between each sample. The Bray-Curtis dissimilarity metric compares two communities based on the number or relative abundance of each taxon present in at least one of the communities. When we calculated these values, we assumed that the set of dissimilarities calculated across a group was independent, even when the same child was paired to other children multiple times. These distance matrices were used for principal coordinates analysis (PCoA) to create ordinations. The two principal components that explained the most variation were used to create biplots.

